# Structural Analysis of Inhibitor Binding to CAMKK1 Identifies Features Necessary for Design of Specific Inhibitors

**DOI:** 10.1101/292797

**Authors:** André da Silva Santiago, Rafael M. Couñago, Priscila Zonzini Ramos, Paulo H. C. Godoi, Katlin B. Massirer, Opher Gileadi, Jonathan M. Elkins

## Abstract

The calcium/calmodulin-dependent protein kinases (CAMKKs) are upstream activators of CAMK1 and CAMK4 signalling and have important functions in neural development, maintenance and signalling, as well as in other aspects of biology such as Ca^2+^ signalling in the cardiovascular system. To support the development of specific inhibitors of CAMKKs we have determined the crystal structure of CAMKK1 with two ATP-competitive inhibitors. The structures reveal small but exploitable differences between CAMKK1 and CAMKK2, despite the high sequence identity, which could be used in the generation of specific inhibitors. Screening of a kinase inhibitor library revealed molecules that bind potently to CAMKK1. Isothermal titration calorimetry revealed that the most potent inhibitors had binding energies largely dependent on favourable enthalpy. Together, the data provide a foundation for future inhibitor development activities.

## Introduction

The calcium/calmodulin-dependent kinases (CAMKs) are of great interest due to their important functions in calcium signalling and especially in neuronal development^1–3^. The CAMK family is formed by CAMK1, CAMK2, CAMK3 (eEF2 kinase), CAMK4 and the CAMK kinases (CAMKKs)^1,4^. Each member of the CAMK family is composed of a catalytic protein kinase domain followed by an auto-inhibitory sequence, which is partially overlapped with a calmodulin-binding domain (CBD). The auto-inhibitory sequence keeps the CAMKs in an inactive state until activation by calcium/calmodulin (Ca^2+^/CaM). Activation occurs by Ca^2+^/CaM binding to the CBD of a CAMK, which releases the auto-inhibitory sequence from the kinase domain^1,2^.

CAMK1, CAMK4 and CAMKKs have a pivotal role in the cascade responsible for Ca^2+^/CaM signalling. This cascade is known to be crucial for neuronal maturation and axon elongation^4^. The CAMKKs are Ser/Thr kinases and the upstream activator kinases of the CaM-stimulated kinase cascade in which activation of CAMK1 and CAMK4 requires phosphorylation at Thr177 and Thr196, respectively, by CAMKKs. There are two CAMKKs, CAMKK1 (CAMKKα) and CAMKK2 (CAMKKβ)^5,6^. There is extensive data supporting the importance of CAMKKs in neurite development, axon elongation, long term potentiation, and long-term memory (LTM), via activation of cAMP-responsive element binding protein (CREB), which is essential for LTM formation^7,8^. Together the data support the value of investigating the roles of CAMKKs in neurodegenerative diseases^1,9^. Furthermore, given the importance of Ca^2+^ signalling in cardiac function it is not surprising that there are important roles for CAMKKs in cardiovascular biology. For example, CAMKK1 was recently discovered to be involved in the regulation of the mesenchymal stem cell secretome^10^.

Given the partially overlapping roles of CAMKK1 and CAMKK2, the commonality of some substrate proteins (such as CAMK1 and CAMK4) and possibility of cross-regulation of expression levels it is a challenge to assign specific roles to CAMKK1 or CAMKK2. A specific inhibitor of either protein would be a valuable research tool for this purpose. So far, the only available CAMKK inhibitors are not specific for one CAMKK protein or indeed over other human kinases. For example, it has been demonstrated that STO-609^11^, the most commonly used inhibitor of CAMKKs, is able to bind to at least seven different kinases with potency similar to CAMKK1 (ERK8, MNK1, AMPK, CK2, DYRK2, DYRK3, and HIPK2)^12^. To aid the development of CAMKK1-or CAMKK2-specific inhibitors we have screened a library of kinase inhibitors against CAMKK1 and determined the X-ray crystal structure of CAMKK1 kinase domain in complex with two different inhibitors. Despite the high sequence similarity between CAMKK1 and CAMKK2 (70% over the kinase domain), comparison of the inhibitor-bound CAMKK1 structures with the existing structures of CAMKK2 reveal small but exploitable differences between the ATP-binding sites that might be used for the future design of specific inhibitors.

## Results

### Identification of kinase inhibitors that are potent inhibitors of CAMKK1

Highly pure CAMKK1 kinase domain protein (residues Gln124-Lys411, hereafter called CAMKK1-KD) was successfully produced using an *E. coli* expression system. This protein was used to screen a library of 378 kinase inhibitors, which identified 86 molecules that bound to CAMKK1 causing a change in protein melting temperature (ΔT_m_) of greater than 2 °C (see full list in Supplementary Information). Previously, two large scale kinase screening efforts published by Davis *et al.*^13^ and Anastassiadis *et al*^14^ included 9 and 6 (respectively) out of the 86 inhibitors identified here. Although there are only 9 inhibitors in common, the dissociation constant (*K*_D_) values measured by Davis *et al.* nevertheless show a correlation (*R*^2^ = 0.52) with the ΔT_m_ values (Figure 1A) allowing us to estimate *K*_D_ for all of the inhibitors identified in our assay. Approximately, a ΔT^m^ of > 7 °C corresponds to a *K*_D_ < 100 nM. Binding curves for inhibitors with ΔT_m_ of > 7 °C are shown in Figure 1B, and the top 20 most potent inhibitors of CAMKK1 are shown in Table 2.

**Figure 1.**
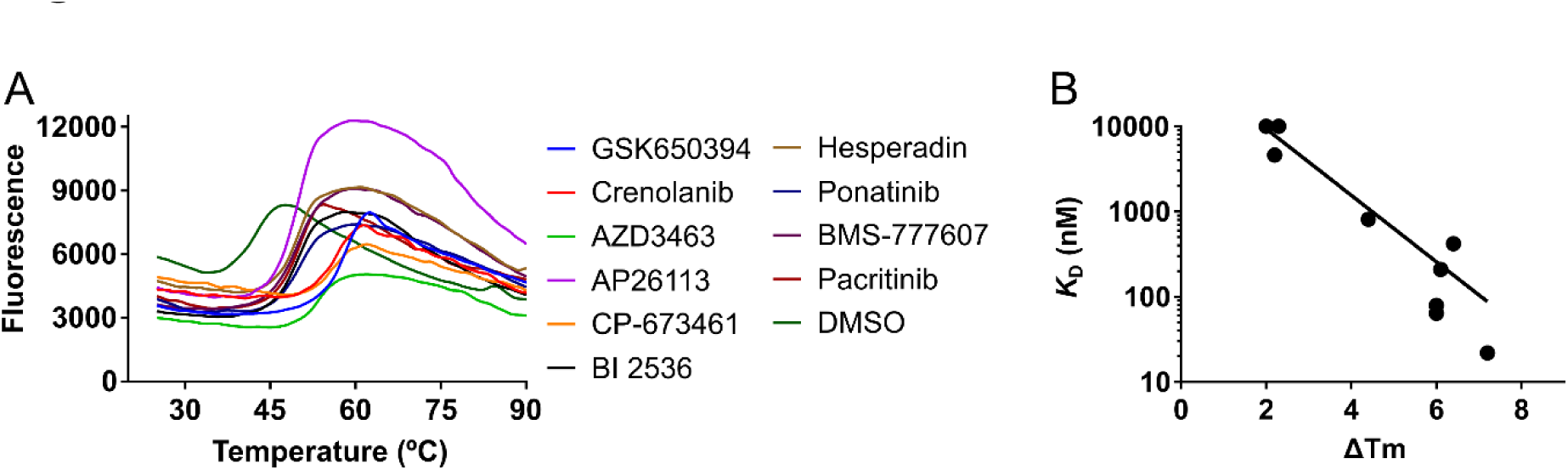
Compound screening identified various inhibitors of CAMKK1. **A.** Thermal melting curves for CAMKK1 protein in the presence of various compounds or DMSO as a reference. **B.** Correlation between change in melting temperature and the dissociation constant *K*_D_ for nine inhibitors where *K*_D_ values were measured by Davis *et al.*^13^. The ΔT_m_ and *K*_D_ values for these nine inhibitors can be found in the Supplementary Information.

**Table 2.**
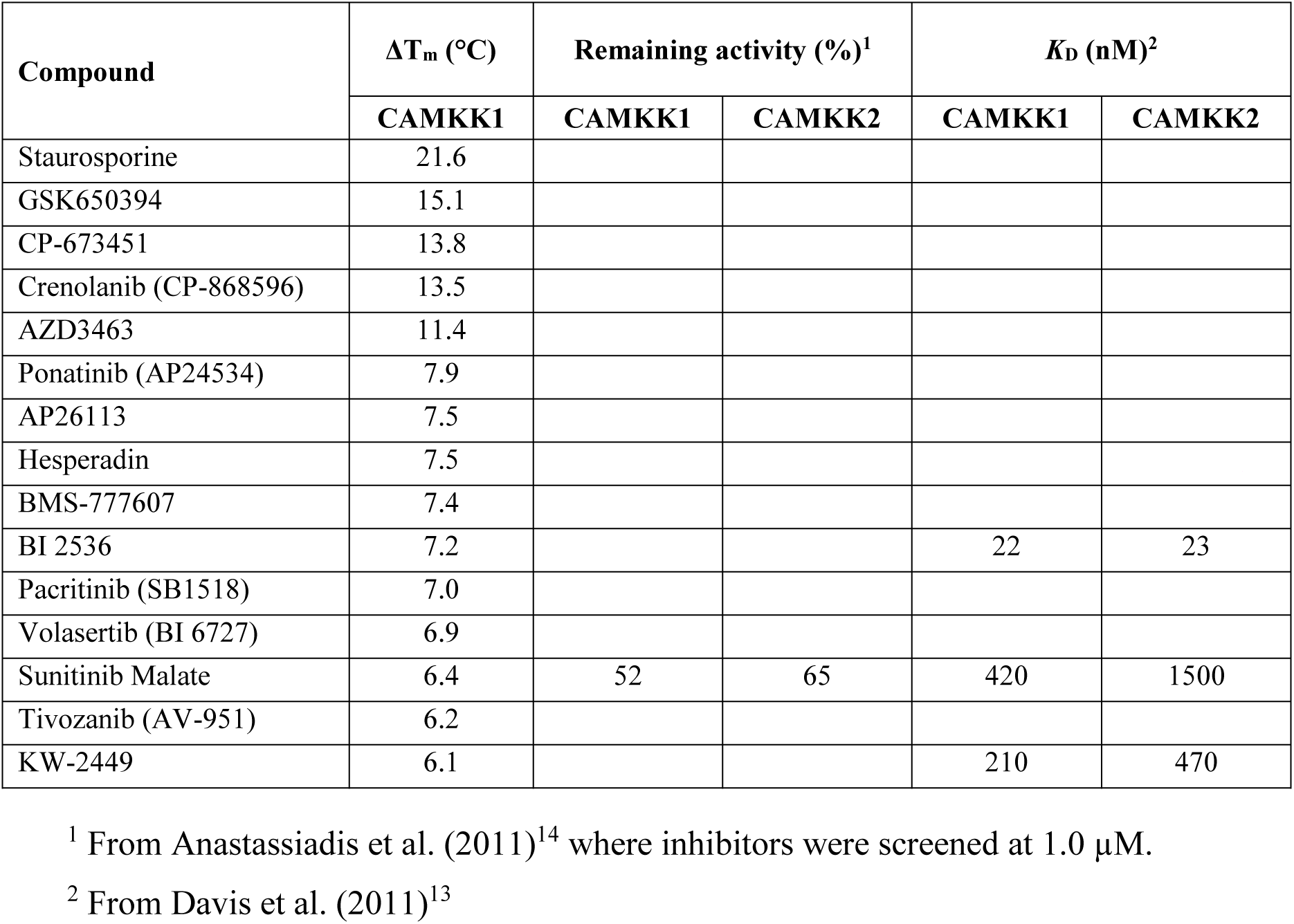
Top hits from DSF screening of a kinase inhibitor library against CAMKK1. All data can be found in the Supplemental Information. Where percentage remaining activity or *K*_D_ values are available in the screening data published by Anastassiadis et al.^14^ or Davis et al.^13^ they are included in the table. Additional activity or *K*_D_ values are found in the Supplemental Information.

| Compound | $\Delta T_m$ (°C) | Remaining activity (%)<sup>1</sup> | | $K_D$ (nM)<sup>2</sup> | |
| --- | --- | --- | --- | --- | --- |
|  | CAMKK1 | CAMKK1 | CAMKK2 | CAMKK1 | CAMKK2 |
| Staurosporine | 21.6 |  |  |  |  |
| GSK650394 | 15.1 |  |  |  |  |
| CP-673451 | 13.8 |  |  |  |  |
| Crenolanib (CP-868596) | 13.5 |  |  |  |  |
| AZD3463 | 11.4 |  |  |  |  |
| Ponatinib (AP24534) | 7.9 |  |  |  |  |
| AP26113 | 7.5 |  |  |  |  |
| Hesperadin | 7.5 |  |  |  |  |
| BMS-777607 | 7.4 |  |  |  |  |
| BI 2536 | 7.2 |  |  | 22 | 23 |
| Pacritinib (SB1518) | 7.0 |  |  |  |  |
| Volasertib (BI 6727) | 6.9 |  |  |  |  |
| Sunitinib Malate | 6.4 | 52 | 65 | 420 | 1500 |
| Tivozanib (AV-951) | 6.2 |  |  |  |  |
| KW-2449 | 6.1 |  |  | 210 | 470 |

<sup>1</sup> From Anastassiadis et al. (2011)<sup>14</sup> where inhibitors were screened at 1.0 μM.
<sup>2</sup> From Davis et al. (2011)<sup>13</sup>

Among the compounds identified as potent inhibitors of CAMKK1 are the ALK/IGF1R inhibitor AZD3463, the PDGFR inhibitors CP-673451, the PDGFR/FLT3 inhibitor Crenolanib, and the SGK inhibitor GSK650394^15^ (which we previously also identified as a CAMKK2 inhibitor). The range of different chemotypes identified as potent CAMKK1 inhibitors suggests CAMKK1 as a common off-target of existing kinase inhibitors, and that development of CAMKK1-specific inhibitors will require a design that incorporates CAMKK1-specific structural features.

### CAMKK1 has a similar overall structure to CAMKK2

Crystallisation trials of CAMKK1-KD with a selection of the strongest binding inhibitors from the DSF analysis yielded crystals in complex with Crenolanib, AZD3463, AP26113, hesperadin and GSK650394. Crystals were also obtained with STO-609 which is well-known as a CAMKK inhibitor^9,11^. However, crystals that diffracted X-rays to high resolution were only obtained with hesperadin and GSK650394. From these crystals it was possible to determine the crystal structure of CAMKK1 to 2.1 Å resolution in complex with hesperadin and 2.2 Å resolution in complex with GSK650394 (Table 1). The two crystal structures were in different crystal forms, with two or four CAMKK1 molecules in the asymmetric unit. The final model for the co-crystal structure with hesperadin consists of residues 124-163 and 193-411, one molecule of hesperadin per CAMKK1 monomer and 271 water molecules. The disordered residues from 164-192 that are not visible in the structure are the RP-insert domain, which is implicated in substrate recognition. This RP-insert domain was also disordered in the co-crystal structure with GSK650394.

**Table 1.** Data collection and refinement statistics

| Ligand | Hesperadin | GSK650394 |
| --- | --- | --- |
| PDB ID | 6CCF | 6CD6 |
| **Data collection** |  |  |
| X-ray source | APS 24-ID-E | APS 24-ID-E |
| Wavelength (Å) | 0.97918 | 0.97918 |
| Space group | *P*2<sub>1</sub> | *P*2<sub>1</sub> |
| Cell dimensions<br>*a*, *b*, *c* (Å); $\beta$ (°) | 49.2, 83.2, 80.1; 97.3 | 44.4, 96.0, 141.3; 90.03 |
| Molecules / asymmetric unit | 2 | 4 |
| Resolution (Å)* | 19.87-2.10 (2.16-2.10) | 19.94-2.20 (2.26-2.20) |
| No. of unique reflections* | 37,282 (3,072) | 59,443 (4,575) |
| *R*<sub>merge</sub> (%)* | 11.1 (98.7) | 8.30 (127.8) |
| Mean *I*/σ(*I*) * | 9.8 (1.7) | 12.1 (1.5) |
| Mean CC(1/2)* | 0.999 (0.893) | 1.000 (0.500) |
| Completeness (%)* | 99.5 (99.8) | 98.8 (98.3) |
| Redundancy* | 5.2 (4.7) | 7.1 (7.4) |
| **Refinement** |  |  |
| Resolution (Å) | 19.87-2.10 (2.16-2.10) | 19.94-2.20 (2.26-2.20) |
| *R*<sub>cryst</sub> / *R*<sub>free</sub> (%) | 21.1 / 24.5 | 23.4 / 28.5 |
| No. of atoms (protein / solvent / ligand) | 4295 / 272 / 81 | 8323 / 110 / 116 |
| Mean *B*-factors (protein / solvent / ligand)<br>(Å<sup>2</sup>) | 35 / 38 / 39 | 62 / 49 / 45 |
| R.m.s.d. bond lengths (Å), angles (°) | 0.008, 1.28 | 0.009, 1.25 |
| **Ramachandran statistics (%)** |  |  |
| Favored / Allowed / Outliers | 97.3 / 2.7 / 0 | 96 / 3.9 / 0.1 |

\*Values in parentheses represent the highest resolution shell

Overall, the CAMKK1 kinase domain has the expected protein kinase fold with an N-terminal lobe mostly of β-strands connected by a flexible linker (hinge region) to a predominantly α-helical C-terminal lobe (Figure 2). In both the structures, the inhibitors hesperadin and GSK650394 are bound to the ATP-binding pocket located between the N-lobe and C-lobe. As expected from the relatively high sequence identity to CAMKK2 (70% over the kinase domain), the CAMKK1 kinase domain structure can be superimposed with the CAMKK2 kinase domain structure (PDB ID 6BKU) with a low r.m.s.d. of 0.66 Å over 261 Cα atoms.

**Figure 2.**
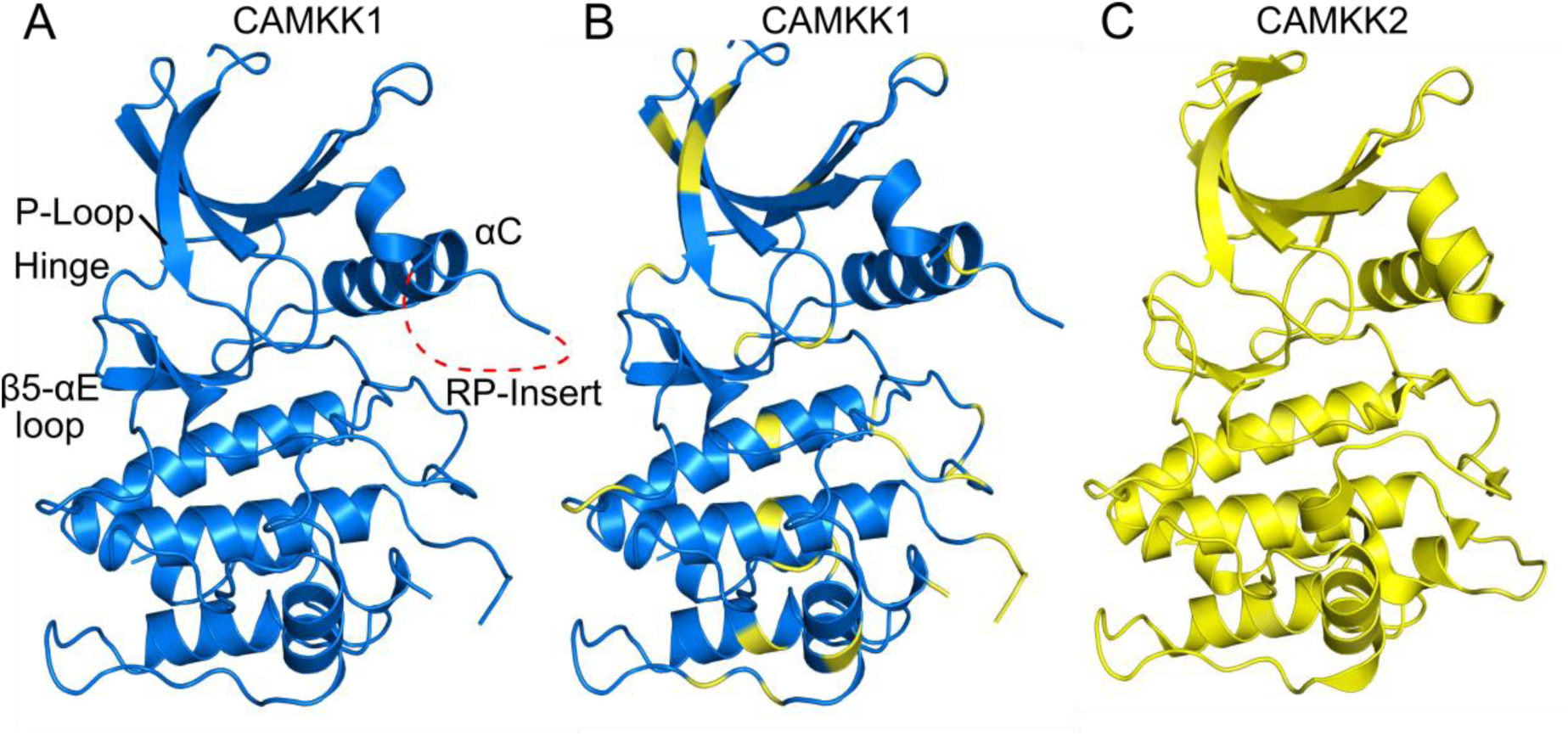
Overall structure of CAMKK1 and comparison to CAMKK2. **A.** CAMKK1 structure. Relevant structural features described in the text are annotated. **B.** CAMKK1 coloured in blue, but with residues not-conserved in CAMKK2 coloured in yellow. **C.** CAMKK2 structure in complex with GSK650394 (PDB ID 6BKU). The structures of CAMKK1 and CAMKK2 with GSK650394 can be superimposed with a r.m.s.d. of 0.66 over 261 Cα atoms.

CAMK1 and CAMK4 have an αD helix (e.g. ^108^LFDRIVEKGF^117^ in CAMK1D, PDB: 2JC6) which is involved in a hydrophobic interaction with the calmodulin binding domain during activation of CAMK1 or CAMK4 by Ca^2+^/CaM ^9^. This helix may also be involved with binding the RP-insert of CAMKKs during activation of the CAMKs ^9^. The positively charged RP-insert from CAMKKs is necessary for selective recognition of the negatively charged αD from CAMK1 and CAMK4^9^. In CAMKK1 (as well as CAMKK2) there is no αD helix (Figure 2); it is replaced by a loop with one proline that is conserved in the CAMKKs (CAMKK1: ^235^KG<u>P</u>VMEVPCDKPF^247^, CAMKK2: ^272^QG<u>P</u>VMEVPTLKPL^284^). Therefore the mechanisms by which CAMKKs interact with calmodulin for activation might be different compared to the mechanism of CAMK1 activation^9^. The calmodulin binding domain in CAMKK1 is at the C-terminus of the protein, comprising the residues ^438^VRLIPSWTTVILVKSMLRK^456^, immediately after the auto-inhibitory domain^16^, and was not included in the construct used for crystallisation.

### Analysis of inhibitor binding to CAMKK1

In both crystal structures the inhibitors were clearly visible in the electron density and could be convincingly modelled (Figure 3, 4). The inhibitors bind as expected in the ATP-binding pocket, between the N-terminal and C-terminal lobes of the kinase domain. In the structure with GSK650394 the inhibitor is bound in a similar manner to that seen in CAMKK2 (PDB ID 6BKU), with two hydrogen bonds to the kinase hinge region, one to the backbone carbonyl atom of Asp231 and the other to the backbone nitrogen atom of Leu233 (Figure 3C). There is a favourable aromatic π-π interaction between the benzoic acid ring of GSK650394 and the side-chain of Phe230, and a hydrophobic interaction between the phenyl ring of GSK650394 and Pro237 on the hinge. The carboxylic acid of GSK650394 forms a hydrogen bond with the side-chain of Lys157, for a total of three hydrogen bonds in direct interaction with CAMKK1. There is also a hydrogen bond to a water molecule which is bound by the side-chain of the αC residue Glu199 and the backbone nitrogen of Phe294 (Figure 3D). ITC measurements showed that CAMKK1 binds GSK650394 strongly, with a *K*_D_ of 4 nM (Figure 3F).

**Figure 3.**
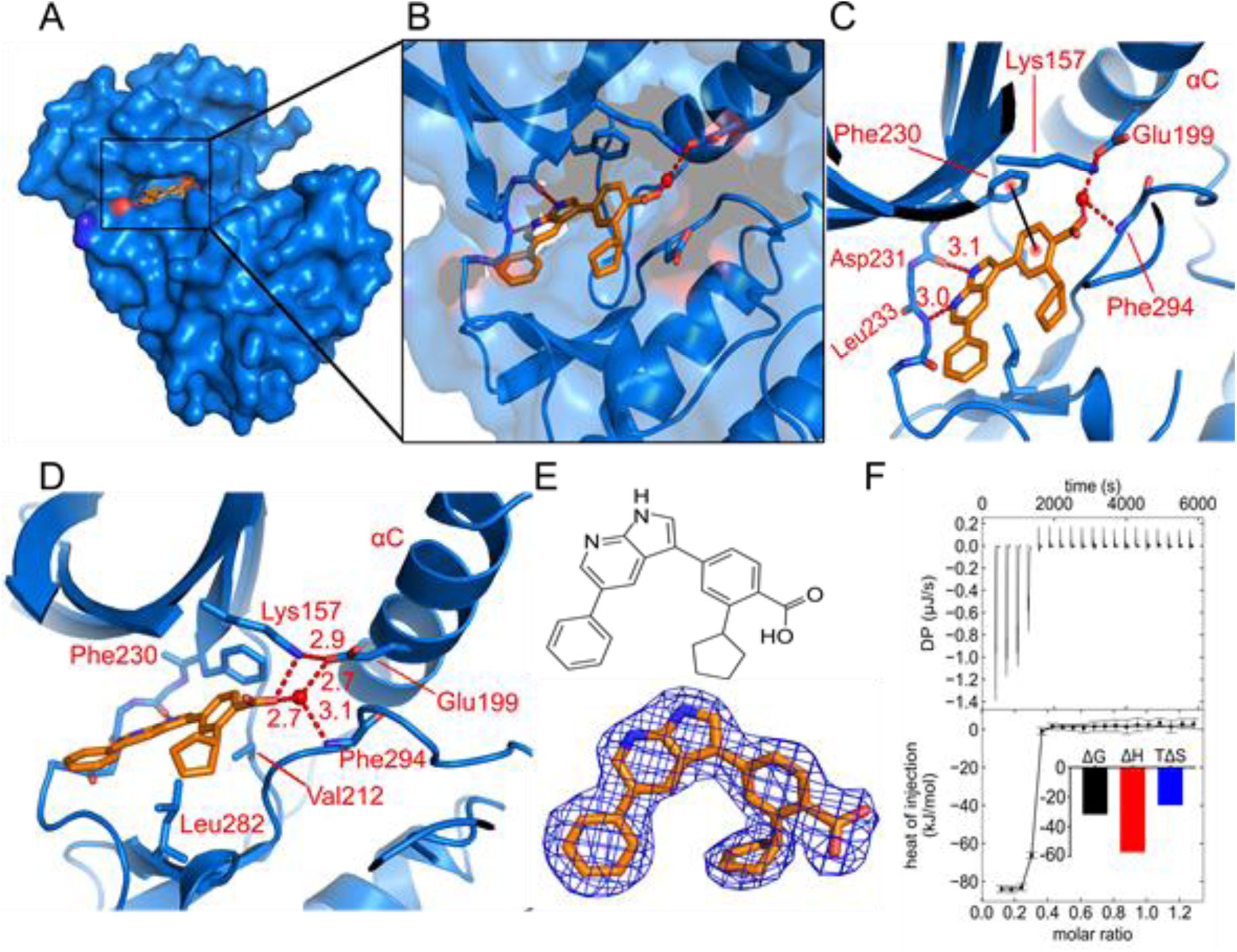
The binding of inhibitor GSK650394 to CAMKK1. **A, B.** GSK650394 (shown in orange) binds in the ATP-binding pocket of CAMKK1 (shown in blue). **C.** GSK650394 binds to the hinge region of CAMKK1 via two hydrogen bonds from its pyrrolo[2,3-b]pyridine moiety to Asp231 and Leu233, as well as a π-stacking interaction between the Phe230 and the benzoic acid moiety. **D.** A water molecule bridges among Glu199, Phe294 and the carboxylic acid group from GSK650394. **E.** Chemical structure of GSK650394, and a 2*F*_o_-*F*_c_ electron density map contoured at 1.0σ for GSK650394 in the crystal structure. **F.** ITC measured a *K*_D_ of 4 nM for the interaction, which had favourable enthalpy and unfavourable entropy of binding.

**Figure 4.**
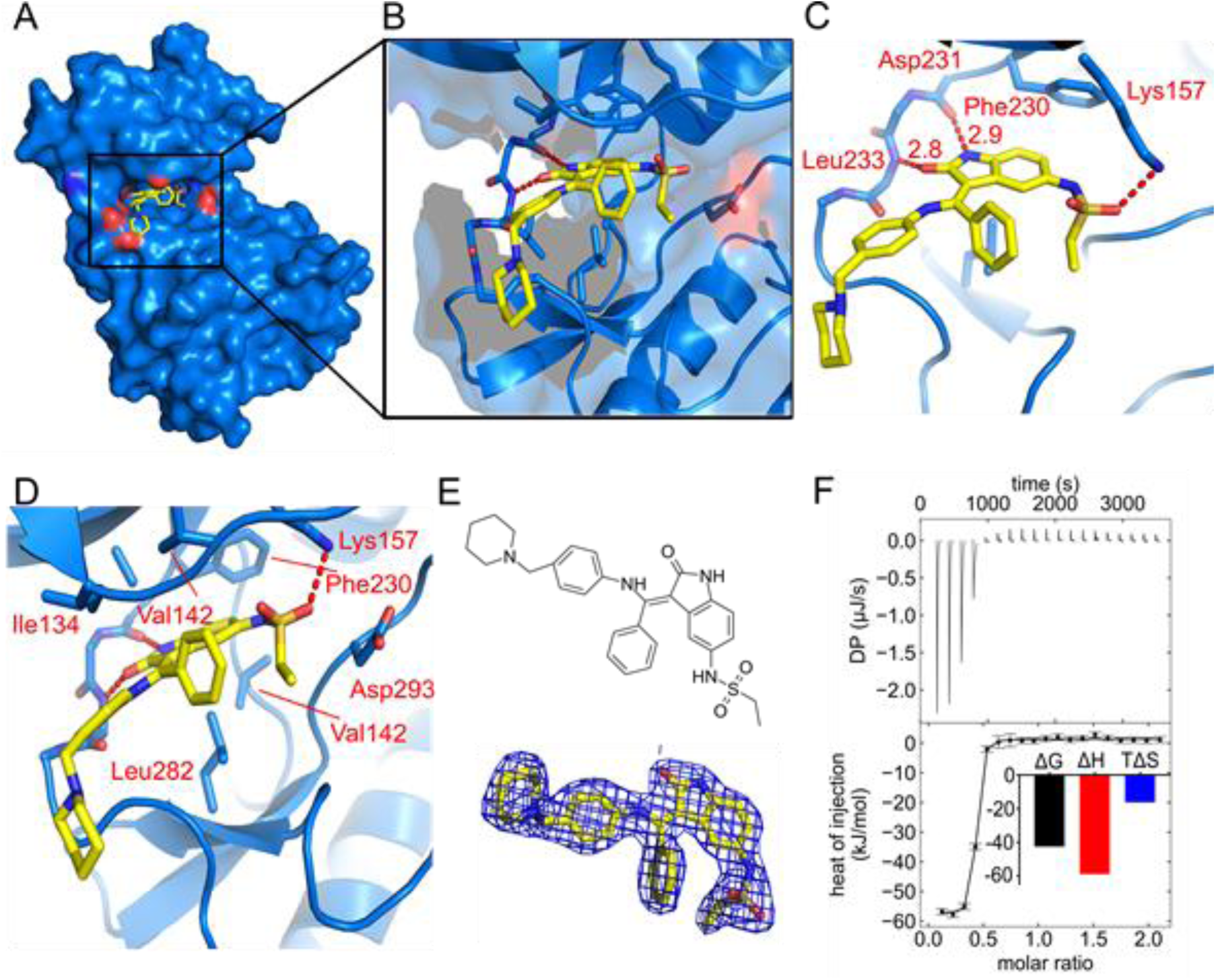
The binding of inhibitor hesperadin to CAMKK1. **A, B.** Hesperadin (shown in yellow) binds in the ATP-binding pocket of CAMKK1 (shown in blue). **C.** Hesperadin binds to the hinge region of CAMKK1 via two hydrogen bonds from its oxindole moiety to Asp231 and Leu233. **D.** A hydrogen bond between a sulfonamide oxygen of hesperadin and Lys157 contributes to the binding energy. **E.** Chemical structure of hesperadin, and 2*F*_o_-*F*_c_ electron density map contoured at 1.0σ for hesperadin in the crystal structure. **F.** ITC measured a *K*_D_ of 26 nM for the interaction, which had favourable enthalpy and unfavourable entropy of binding.

In the structure with hesperadin, the inhibitor again forms two hydrogen bonds to the hinge, via its oxindole moiety to the backbone carbonyl atom of Asp231 and the backbone nitrogen atom of Leu233 (Figure 4C). Hesperadin also forms a hydrogen bond via its sulphonamide to the side-chain of Lys157, while the oxindole ring forms a hydrophobic interaction with Phe230 (Figure 4C, 4D). ITC measurements showed that CAMKK1 binds hesperadin with a *K*_D_ of 26 nM (Figure 4F).

### The most potent CAMKK1 inhibitors bind due to favourable enthalpy

After seeing that hesperadin and GSK650394 were extremely potent inhibitors of CAMKK1, we measured ITC data for four additional inhibitors that bound strongly to CAMKK1 in the DSF assay (Figure 5, Table 3), to confirm if the low dissociation constants and enthalpy-dependent binding seen with hesperadin and GSK650394 were common features of the identified CAMKK1 inhibitors. The data show that all of the tested inhibitors are extremely potent binders of CAMKK1, and furthermore in all cases the binding is heavily enthalpy-dependent.

**Figure 5.**
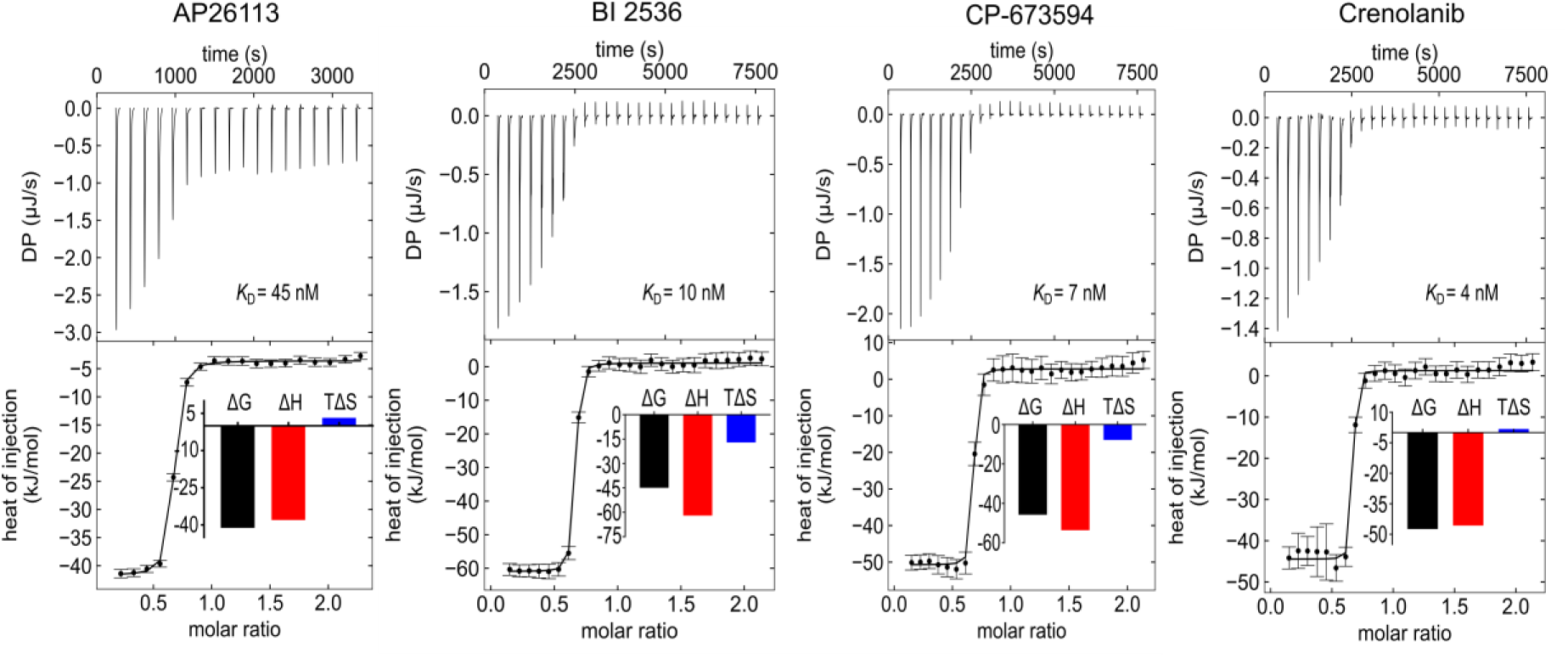
Dissociation constant measurements by ITC for four additional inhibitors that promoted a change in the melting temperature (ΔT_m_) of CAMKK1 by more than 7 ºC. The top part of each figure shows the injection heats, the bottom part shows the fitted binding isotherms (a single-site model was used), and the insets show the binding energies in kcal/mol.

**Table 3.** Binding affinities for selected inhibitors measured by ITC at 20 °C, showing the good correlation with thermal stabilisation of CAMKK1 measured by DSF.

| Inhibitor | $K_D$ (nM) | $\Delta H$<br>(kcal/mol) | Fraction<br>incompetent A<br>(ligand in cell) | Fraction<br>incompetent B<br>(protein) | $\Delta T_m$<br>(°C) |
| --- | --- | --- | --- | --- | --- |
| GSK650394 | 4 | -20.5 | 0.7 | 0.0 | 15.1 |
| Crenolanib | 4 | -10.9 | 0.4 | 0.0 | 13.5 |
| CP-673451 | 7 | -12.8 | 0.4 | 0.0 | 13.8 |
| BI 2536 | 10 | -14.8 | 0.4 | 0.0 | 7.2 |
| Hesperadin | 26 | -14.1 | 0.6 | 0.0 | 7.5 |
| AP26113 | 45 | -9.1 | 0.4 | 0.0 | 7.5 |

### Two active site differences could allow the design of specific inhibitors

The ATP-binding sites of CAMKK1 and CAMKK2 are highly conserved, which is reflected in the similarity of the inhibitor binding profiles of CAMKK1 and CAMKK2 seen previously^13,14^ Comparison of the CAMKK1 structures to CAMKK2 reveals two possible options for the design of specific inhibitors (Figure 6). The first is that there are differences in the hydrophobic “back-pocket” region around the kinase regulatory spine (R-spine). In particular, Leu228 in CAMKK1 is Met265 in CAMKK2 and this difference would presumably affect the movement of the R-spine residues in response to inhibitors that disrupt the R-spine, i.e. type II kinase inhibitors. A type II inhibitor would have the potential to gain selectivity between CAMKK1 and CAMKK2 by taking advantage of this and other non-conserved residues outside the ATP-binding pocket itself.

**Figure 6.**
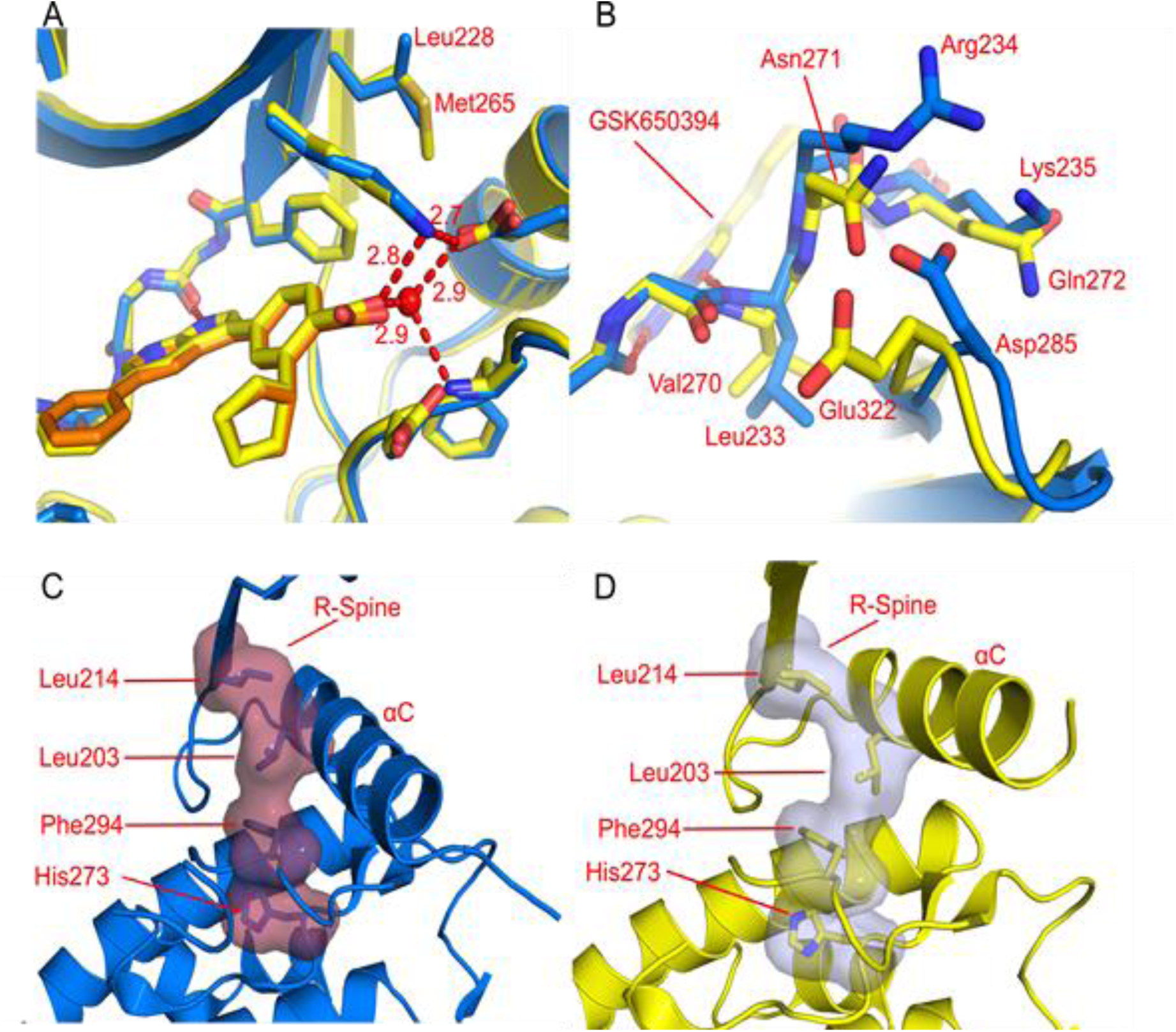
Structural differences between CAMKK1 (shown in blue) and CAMKK2 (shown in yellow) that could be exploited to design specific inhibitors. **A.** Adjacent to the kinase R-spine Leu228 in CAMKK1 is Met265 in CAMKK2. An inhibitor that disrupts the R-spine (a type 2 inhibitor) could engage this amino acid **B.** Due to different interactions between the hinge region (Arg234 in CAMKK1) and the β4-β5 loop (Asp285 in CAMKK1) the hinge region is significantly further away from the ATP-binding site in CAMKK2 (1.2 Å). This could be exploited by an inhibitor that interacts with the backbone of Arg234 (or Asp285 for a CAMKK2 inhibitor). **C.** View of the R-spine residues and αC helix of CAMKK1 when bound to GSK650394. **D.** View from the same angle as **C** of the R-spine residues and αC helix of CAMKK1 when bound to hesperadin, showing the deformation of the R-spine and the more outward position of αC in the presence of this inhibitor, and the flexibility of these residues.

However, a more significant difference between CAMKK1 and CAMKK2 is seen in the kinase hinge region (Figure 6B). The hinge residues Leu233, Arg234 and Lys235 are replaced in CAMKK2 by Val270, Asn271 and Gln272. Although none of these amino acid side-chains are directed into the ATP-binding site, these sequence differences cause a significant difference in the conformation of the protein backbone in this region, greater than 1 Å difference in position. Especially important is CAMKK1 Leu233, which is larger than the equivalent valine residue in CAMKK2 and to be accommodated in the same space requires the different backbone conformation observed. The differences in the hinge residues Arg234 and Lys235 which interact with the β6-β7 loop are also likely to contribute to the different conformation, especially since the key residue on the β6-β7 loop, Asp285, is also not conserved in CAMKK2.

## Discussion

The compound screening data showed that a significant number of kinase inhibitors interact with CAMKK1, despite none of these molecules being originally designed as a CAMKK inhibitor. This suggests that CAMKK1 is an easily inhibited kinase, and given the strength of binding observed, that alteration of Ca^2+^/CaM signalling may be a significant cause of some of the phenotypes observed with some of these inhibitors. It also suggests that achieving high potency with a newly designed CAMKK1-specific inhibitor will be possible, especially given the relatively high *K*_M_(ATP) previously reported for CAMKK^16^,^17^.

As the ATP-binding sites of CAMKK1 and CAMKK2 are so similar, for future design of a specific inhibitor there are two possibilities: an allosteric inhibitor, or an inhibitor which can take advantage of the small differences that do exist to achieve specificity. The two different crystal structures of CAMKK1 with different inhibitors show a large difference in the position of helix αC (Figure 6C, 6D). In the structure with hesperadin αC is in an inactive αC-out conformation, with a disrupted kinase R-spine (Figure 6D). The R-spine is a set of four residues in kinases that line up to form a hydrophobic spine when a protein kinase is in an active conformation^18^. Since it appears that hesperadin does not directly push out the αC helix, this indicates flexibility of this region of the protein and therefore that a type II inhibitor (or indeed an allosteric, type III, inhibitor) binding to and disrupting the R-spine would be possible. The inhibitor screening data shows that a number of known type II inhibitors such as ponatinib bind strongly to CAMKK1. Given that there are sequence differences between CAMKK1 and CAMKK2 in this region of the R-spine and αC, this could be a way to achieve selectivity. Finally, the significant differences in the conformation of the C-terminal part of the kinase hinge region (residues Arg234-Gly236, Figure 6B) could be exploited by suitably-designed type I kinase inhibitors.

## Materials and Methods

### DNA constructs

A full length coding DNA clone for human CAMKK1 was obtained from the Mammalian Gene Collection (IMAGE Consortium Clone ID 5751570, corresponding to CAMKK1 isoform A, Uniprot Q8N5S9-1) and used as template for PCR amplification of truncated constructs. A construct comprising the kinase domain (Gln124 to Lys411) was amplified using the primers: forward TACTTCCAATCCATGCAGCTGAACCAGTACAAGCTG, reverse TATCCACCTTTACTGTCACTTGGTCACCCAAGGATGCAACTTGATGTC, and cloned into vector pNIC28-Bsa4 by ligation independent cloning^19^. The construct was verified by DNA sequencing.

### Protein expression and purification

CAMKK1-Q124-K411 (CAMKK1-KD) in pNIC28-Bsa4 was transformed into chemically competent *Escherichia coli* BL21(DE3)-R3 cells that co-express λ-phosphatase and three rare tRNAs (plasmid pACYC-LIC+). Colonies were used to inoculate 50 mL of LB media supplemented with 5 mM betaine, 50 µg/mL kanamycin and 35 µg/mL chloramphenicol which was left shaking at 37 °C overnight. This culture was used to inoculate 1.5 L of Terrific Broth containing 50 µg/mL kanamycin at 37 °C until the OD_600_ reached 3. The culture was cooled to 18 °C, 0.2 mM of Isopropyl β-D-1-thiogalactopyranoside (IPTG) was added, and left overnight. Cells were collected by centrifugation for 15 min at 7,500 × g at room temperature. The cell pellet was weighed and resuspended in the same volume of 2× binding buffer (1× binding buffer was 50 mM HEPES pH 7.5, 500 mM NaCl, 10% glycerol, 10 mM imidazole, 1 mM tris(2-carboxyethyl)phosphine (TCEP) with protease inhibitors set II (Calbiochem) at 1:200 ratio) and frozen at -80 °C until use.

For purification, the cell pellet was thawed and sonicated on ice. Polyethyleneimine (pH 7.5) was added to the lysate to a final concentration of 0.15% (w/v) and the sample was centrifuged at 53,000 × g for 45 min at 4 °C. The supernatant was loaded onto an IMAC column (5 mL HisTrap FF Crude) and washed in binding buffer containing 30 mM imidazole. The protein was eluted with elution buffer (binding buffer with 300 mM imidazole). To remove the 6xHis-tag, the protein was incubated with TEV protease during dialysis overnight at 4 ºC into gel filtration buffer (20 mM HEPES, 500 mM NaCl, 5% glycerol, 1 mM TCEP, 5% [v/v] glycerol). Protein was further purified using reverse affinity chromatography on Ni-Sepharose followed by gel filtration (Superdex 200 16/60, GE Healthcare). Protein in gel filtration buffer was concentrated to 10.4 mg/mL (measured by UV absorbance with a NanoDrop spectrophotometer (Thermo Scientific) using the calculated molecular weight and estimated extinction coefficient) using 10 kDa molecular weight cut-off centrifugal concentrators (Millipore) at 4 °C.

### Differential scanning fluorimetry

CAMKK1-KD protein was screened against a library of 378 structurally diverse and cell-permeable ATP-competitive kinase inhibitors available from Selleckchem (Houston, TX, United States; catalog No. L1200). DSF measurements were made in a 384-well plate. Each well contained 20 μL of 1 μM kinase in 100 mM potassium phosphate pH 7.0, 150 mM NaCl, 10% glycerol and the Applied Biosystems Protein Thermal Shift dye at the recommended concentration of 1:1000.

The compounds, previously solubilized in DMSO, were used at 10 µM final concentration and 0.1% DMSO. Plates were sealed using optically clear films and transferred to a QuantStudio 6 qPCR instrument (Applied Biosystems). The fluorescence intensity was measured during a temperature gradient from 25 to 95 ºC at a constant rate of 0.05 °C/s and protein melting temperatures were calculated based on a Boltzmann function fitting to experimental data, as implemented in the Protein Thermal Shift Software (Applied Biosystems). Protein in 0.1% DMSO was used as a reference. Compounds that caused a shift in melting temperature of the protein (ΔT_m_) of 2 °C or higher compared to the reference were considered positive hits.

### Crystallisation, data collection and structure determination

For crystallization, inhibitors previously identified were added to purified CAMKK1-KD at a threefold molar excess and the samples were incubated on ice for one hour. Initially, 40 μL of purified protein at 10.4 mg/mL was diluted in 360 μL of gel filtration buffer, passed to a fresh 1.5 mL tube containing 5.8 μL of the inhibitor (10 mM in DMSO), and incubated for one hour. The mixture was concentrated at 4 ºC in an Amicon Ultra 10 kDa MW cutoff concentrator (Millipore) for DMSO removal. Twice, 300 μL of gel filtration buffer was added and the sample concentrated again to 40 μL final volume. Crystallization was performed in SWISSCI 3-well low profile crystallization plates at 6 ºC. Crystals of CAMKK1-KD with hesperadin or GSK650394 grew from a mixture of 100 nL of protein:inhibitor solution and 100 nL of a reservoir solution of 33% PEG2000 MME, 0.1 M potassium thiocyanate, CHC buffer (0.1 M each citric acid, HEPES, and CHES) pH 7.5. Crystals were cryo-protected in reservoir solution supplemented with 30% ethylene glycol before flash-freezing in liquid nitrogen for data collection (Table1).

The data was integrated with XDS^20^ and scaled using AIMLESS from the CCP4 software suite^21^. The structure was solved by molecular replacement using Phaser^22^ and the kinase domain of CAMKK2 as the search model (PDB ID 2ZV2)^9^. Refinement was performed using REFMAC5^23^ and Coot^24^ was used for model building. Structure validation was performed using MolProbity^25^.

### Isothermal calorimetry

Measurements were made using a MicroCal AutoiITC200 (Malvern, United Kingdom), at 20 ºC with 1000 rpm stirring. For all measurements CAMKK1-KD protein was dialysed overnight against gel filtration buffer, and the dialysis buffer was used to dilute the inhibitors. CAMKK1-KD protein was titrated into a solution containing the inhibitor. The concentrations used for each measurement were: hesperadin 27 µM, CAMKK1-KD 270 µM (measurement made using 2 µL injections and 180 s between each injection); AP26113, BI2536, Crenolanib or CP-673451 26.7 µM, CAMKK1-KD 267 µM (measurements made using 1.5 µL injections and 300 s between each injection); GSK650394 30.8 µM, CAMKK1-KD 247 µM (measurement made using 1.5 µL injections and 300 s spacing between each sample). ITC data was analysed with NITPIC, and SEDPHAT; figures were made using GUSSI^26^.

### Data availability statement

The coordinates and structure factors for the crystal structures of CAMKK1 have been deposited in the Protein Data Bank with accession codes 6CCF and 6CD6.

## Acknowledgements

This work is based upon research conducted at the Northeastern Collaborative Access Team beamlines (GU51510, GU56413), which are funded by the National Institute of General Medical Sciences from the National Institutes of Health (P41 GM103403). The Pilatus 6M detector on 24-ID-C beam line is funded by a NIH-ORIP HEI grant (S10 RR029205). This research used resources of the Advanced Photon Source, a U.S. Department of Energy (DOE) Office of Science User Facility operated for the DOE Office of Science by Argonne National Laboratory under Contract No. DE-AC02-06CH11357.

The SGC is a registered charity (number 1097737) that receives funds from AbbVie, Bayer Pharma AG, Boehringer Ingelheim, Canada Foundation for Innovation, Eshelman Institute for Innovation, Genome Canada, Innovative Medicines Initiative (EU/EFPIA) [ULTRA-DD grant no. 115766], Janssen, Merck KGaA Darmstadt Germany, MSD, Novartis Pharma AG, Ontario Ministry of Economic Development and Innovation, Pfizer, São Paulo Research Foundation-FAPESP (2013/50724-5), Takeda, and Wellcome [106169/ZZ14/Z]. André da Silva Santiago received a CNPQ-PVE fellowship (152254/2016-1).

## Author contributions

ADSS participated in all parts of the project. RMC collected diffraction data and helped with structure refinement. PZR and OG designed constructs and did cloning. PHCG performed compound screening and data analysis. KBM, OG and JME coordinated the project. ADSS and JME wrote the manuscript. All authors revised the manuscript.

## References

1. Takemoto-Kimura, S. et al. Calmodulin kinases: essential regulators in health and disease. J. Neurochem. 141, 808–818 (2017).

2. Soderling, T. R. The Ca-calmodulin-dependent protein kinase cascade. Trends Biochem. Sci. 24, 232–236 (1999).

3. Takano, T. et al. Discovery of long-range inhibitory signaling to ensure single axon formation. Nat. Commun. 8, 1–17 (2017).

4. Marcelo, K. L., Means, A. R. & York, B. The Ca2+/Calmodulin/CaMKK2 Axis: Nature’s Metabolic CaMshaft. Trends Endocrinol. Metab. 27, 706–718 (2016).

5. Marcelo, K. L. et al. Research Resource: Roles for Calcium/Calmodulin-Dependent Protein Kinase Kinase 2 (CaMKK2) in Systems Metabolism. Mol. Endocrinol. 30, 557–572 (2016).

6. Okuno, S., Kitani, T. & Fujisawa, H. Regulation of Ca(2+)/calmodulin-dependent protein kinase kinase alpha by cAMP-dependent protein kinase: I. Biochemical analysis. J. Biochem. 130, 503–13 (2001).

7. Wayman, G. a, Lee, Y., Tokumitsu, H., Silva, A. & R, T. NIH Public Access. Signal Transduct. 59, 914–931 (2009).

8. Kaitsuka, T. et al. Forebrain-specific constitutively active CaMKKα transgenic mice show deficits in hippocampus-dependent long-term memory. Neurobiol. Learn. Mem. 96, 238–247 (2011).

9. Kukimoto-Niino, M. et al. Crystal structure of the Ca2+/calmodulin-dependent protein kinase kinase in complex with the inhibitor STO-609. J. Biol. Chem. 286, 22570–22579 (2011).

10. Dong, F. et al. A Novel Role for CAMKK1 in the Regulation of the Mesenchymal Stem Cell Secretome. Stem Cells Transl. Med. 6, 1759–1766 (2017).

11. Tokumitsu, H. et al. STO-609, a specific inhibitor of the CA2+/calmodulin-dependent protein kinase kinase. J. Biol. Chem. 277, 15813–15818 (2002).

12. Bain, J. et al. The selectivity of protein kinase inhibitors: a further update. Biochem. J. 408, 297–315 (2007).

13. Davis, M. I. et al. Comprehensive analysis of kinase inhibitor selectivity. Nat. Biotechnol. 29, 1046–1051 (2011).

14. Anastassiadis, T., Deacon, S. W., Devarajan, K., Ma, H. & Peterson, J. R. Comprehensive assay of kinase catalytic activity reveals features of kinase inhibitor selectivity. Nat. Biotechnol. 29, 1039–1045 (2011).

15. Sherk, A. B. et al. Development of a small molecule serum and glucocorticoid-regulated kinase 1 antagonist and its evaluation as a prostate cancer therapeutic. Cancer Res. 2008 68, 7475–7483 (2009).

16. Tokumitsu, H., Wayman, G. a, Muramatsu, M. & Soderling, T. R. Calcium/calmodulin-dependent protein kinase kinase: identification of regulatory domains. Biochemistry 36, 12823–12827 (1997).

17. Kitani, T., Okuno, S. & Fujisawa, H. Regulation of Cat+/Calmodulin-Dependent Protein Kinase Kinase a by cAMP-Dependent Protein Kinase: II. Mutational Analysis1. J Biochem 130, 515–525 (2001).

18. Kornev, A. P., Taylor, S. S. & Ten Eyck, L. F. A helix scaffold for the assembly of active protein kinases. Proc. Natl. Acad. Sci. 105, 14377–14382 (2008).

19. Savitsky, P. et al. High-throughput production of human proteins for crystallization: The SGC experience. J. Struct. Biol. 172, 3–13 (2010).

20. Kabsch, W. Xds. Acta Crystallogr. Sect. D Biol. Crystallogr. 66, 125–132 (2010).

21. Winn, M. D. et al. Overview of the CCP4 suite and current developments. Acta Crystallogr. Sect. D Biol. Crystallogr. 67, 235–242 (2011).

22. McCoy, A. J. et al. Phaser crystallographic software. J. Appl. Crystallogr. 40, 658–674 (2007).

23. Murshudov, G. N. et al. REFMAC5 for the refinement of macromolecular crystal structures. Acta Crystallogr. Sect. D Biol. Crystallogr. 67, 355–367 (2011).

24. Emsley, P., Lohkamp, B., Scott, W. G. & Cowtan, K. Features and development of Coot. Acta Crystallogr. Sect. D Biol. Crystallogr. 66, 486–501 (2010).

25. Chen, V. B. et al. MolProbity: All-atom structure validation for macromolecular crystallography. Acta Crystallogr. Sect. D Biol. Crystallogr. 66, 12–21 (2010).

26. Brautigam, C. A., Zhao, H., Vargas, C., Keller, S. & Schuck, P. Integration and global analysis of isothermal titration calorimetry data for studying macromolecular interactions. Nat. Protoc. 11, 882–894 (2016).

